# Hepatic JARID1a ablation disrupts the transcription adaptation to feeding and alters systemic metabolism

**DOI:** 10.1101/499566

**Authors:** Kacee A. DiTacchio, Diana Kalinowska, Anand Saran, Ashley Byrne, Christopher Vollmers, Luciano DiTacchio

## Abstract

The liver is a key regulator of systemic energy homeostasis whose proper function is dependent on the circadian clock. Here, we show that livers deficient in the oscillator component JARID1a exhibit a dysregulation of genes involved in energy metabolism. Importantly, we find that mice that lack hepatic JARID1a have decreased lean body mass, decreased respiratory exchange ratios, faster production of ketones and increased glucose production in response to fasting. Finally, we find that JARID1a loss compromises the response of the hepatic transcriptome to nutrient availability. In all, ablation of hepatic JARID1a disrupts the coordination of hepatic metabolic programs with whole-body consequences.

## Main

The circadian clock is an endogenous timing mechanism that generates ~24-hour behavioral and physiological oscillations that allow organisms to adapt to the changing environment inherent to the day-night cycle. In recent years, the circadian oscillator has emerged as a critical orchestrator of metabolism and energy homeostasis with important implications to human health. Circadian dysfunction due to environmental factors commonly found in modern lifestyles has been linked to weight gain, metabolic syndrome and diabetes (Albrecht, 2012; Bass and Takahashi, 2010; Feng and Lazar, 2012; Green et al., 2008). Critically, one way in which a high-fat, western-style diet promotes imbalance in energy metabolism is through interference of circadian function (Kohsaka et al., 2007; Marcheva et al., 2010). Conversely, improvement of circadian function via feeding schedule manipulation is able to prevent and reverse the deleterious effects of high fat diet in mice (Chaix et al., 2014; Hatori et al., 2012), underscoring the importance of the circadian system in the maintenance of metabolic homeostasis.

At the molecular level, circadian rhythms originate from a cell-autonomous molecular circuit that impinges on physiology largely through transcriptional control. In mammals, these cell-autonomous oscillators are assembled into tissue-level oscillators which generate local rhythms in physiology. In the liver, the local oscillator is critical for normal function, and its disruption is associated with fatty liver, disruption of glucose homeostasis, and diabetes (Feng et al., 2011; Lamia et al., 2008; Tahara and Shibata, 2016). Interestingly, the hepatic clock is required but not sufficient to generate large-scale transcriptional rhythms. Rather, the hepatic circadian transcriptome arises from an interaction between feeding-derived cues and the circadian clock (Vollmers et al., 2009) through mechanisms that are unclear.

We previously identified the JmjC and AT-rich Interacting Domain protein 1a (JARID1a) as a non-redundant, evolutionarily conserved component of the circadian molecular machinery (DiTacchio et al., 2011). Mechanistically, JARID1a acts as a transcriptional co-activator for CLOCK-BMAL1 by inhibition of HDAC activity, acting as a molecular switch that triggers the transition from the repressive to the active phase of the circadian transcriptional cycle, and in its absence the amplitude of circadian oscillations is severely dampened and the period shortened. In addition, JARID1a has also been found to associate with and participate in the regulation by several transcription factors that have mechanistic links to energy metabolism (Benevolenskaya et al., 2005; Chan and Hong, 2001; Hayakawa et al., 2007). These observations, coupled to its role in the clock, led us to assess the role of JARID1a as a novel contributor of circadian regulation of energy metabolism *in vivo*.

We generated a *Jarid1a* liver-specific knockout (*Jarid1a^LKO^*) mouse model to determine whether JARID1a plays a molecular role at the intersection of circadian and systemic metabolic control, without confounding disruptions to behavioral rhythms. As expected, *Jarid1a^LKO^* mice exhibited normal rhythms in activity and feeding, and unaltered total activity and caloric intake (Figure 1A-D). From 10-weeks of age until the end of the study we observed that *Jarid1a^LKO^* mice exhibited a slight, but statistically significant lower body weight than that of control mice (p<0.05, n=20-24 per group) (Figure 1E and F). This difference in body weight was accentuated under a high fat diet (40% kcal from fat), (p<0.002, n=19-25 per group) (Figure 1E). Since the liver plays an important role in maintenance of blood sugar, we performed a pyruvate tolerance test (PTT), which measures blood glucose levels following intraperitoneal injection of pyruvate. Pyruvate is used as a substrate by the liver to produce glucose and release it into the bloodstream through the process of gluconeogenesis. On both standard and high fat diets, *Jarid1a^LKO^* mice exhibited an increased glucose response upon pyruvate challenge (Figure 1F). This indicates that *Jarid1a^LKO^* mice have increased gluconeogenesis; however, for the duration of the current study, the excess production was insufficient to generate an overt diabetic phenotype as no insulin resistance was detected (Fig S2) and only a slightly sustained glucose response could be observed in *Jarid1a^LKO^* mice of the HFD cohort in an intraperitoneal glucose tolerance test (GTT, Figure 1G).

**Figure 1.**
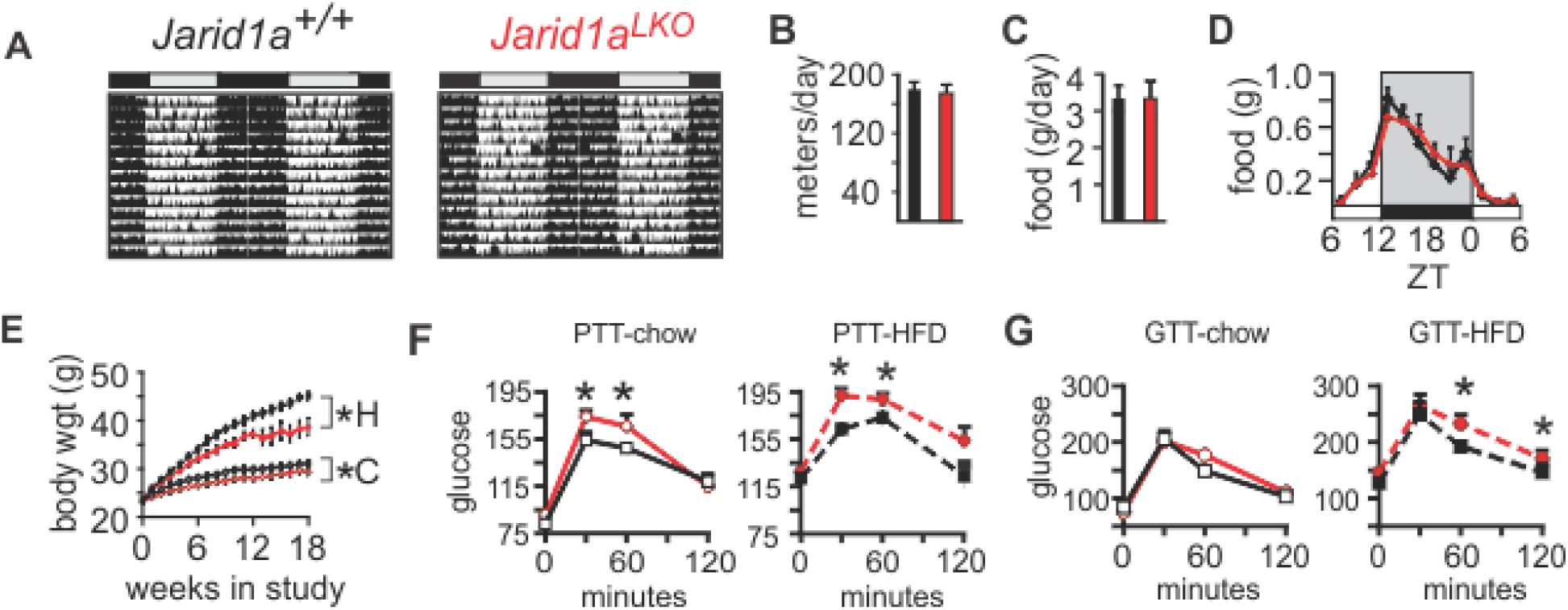
Metabolic phenotype of *Jarid1a^LKO^* mice. (**A**) Representative circadian double-plotted diurnal activity of control and *Jarid1a^LKO^* mice. All actograms obtained are shown in Figure S1. (**B**) Average daily distance traveled by both mouse cohorts (mean+/−s.e.m, n=6/cohort). (**C**) Daily food consumption (mean+/− s.e.m., n=6/cohort) and (**D**) diurnal food consumption profile for control and *Jarid1a^LKO^* mice under 12h:12h light:dark cycle (mean+/− s.e.m. n=6/cohort). (**E**) Weight gain under regular or high-fat (HF) diets (mean+/− s.e.m., control n=19, *Jarid1a^LKO^* n=24). (**F**) Intraperitoneal pyruvate tolerance test (PTT) for mice under regular chow (control n=14, *Jarid1a^LKO^* n=11) or a high-fat diet (HFD; control n=8, *Jarid1a^LKO^* n=11). (**G**) Intraperitoneal glucose tolerance test (GTT) for mice under regular chow (control n=10, *Jarid1a^LKO^* n=10) or a high-fat diet (HFD; control n=14, *Jarid1a^LKO^* n=8). Data for (E-G) are presented as mean+/−s.e.m. * p<0.05, ** p<0.01; p values obtained in (**E**) with repeated measures ANOVA, and (F, G) with permutation tests.

The liver plays a critical role in coordinating energy homeostasis for the whole body during periods of fasting. In additional to glucose production, the liver also catabolizes lipids and produces ketones that can then be used as an alternative energy source. We examined the effects of fasting on blood glucose and ketone levels in *Jarid1a^LKO^* mice versus control animals. We found a slight increasing trend in blood glucose concentration up to 9 hours following food withdrawal in both groups, which may be attributed to an overlap between increased glycogenolysis and gluconeogenesis as fasting progresses (Figure 2A).

**Figure 2.**
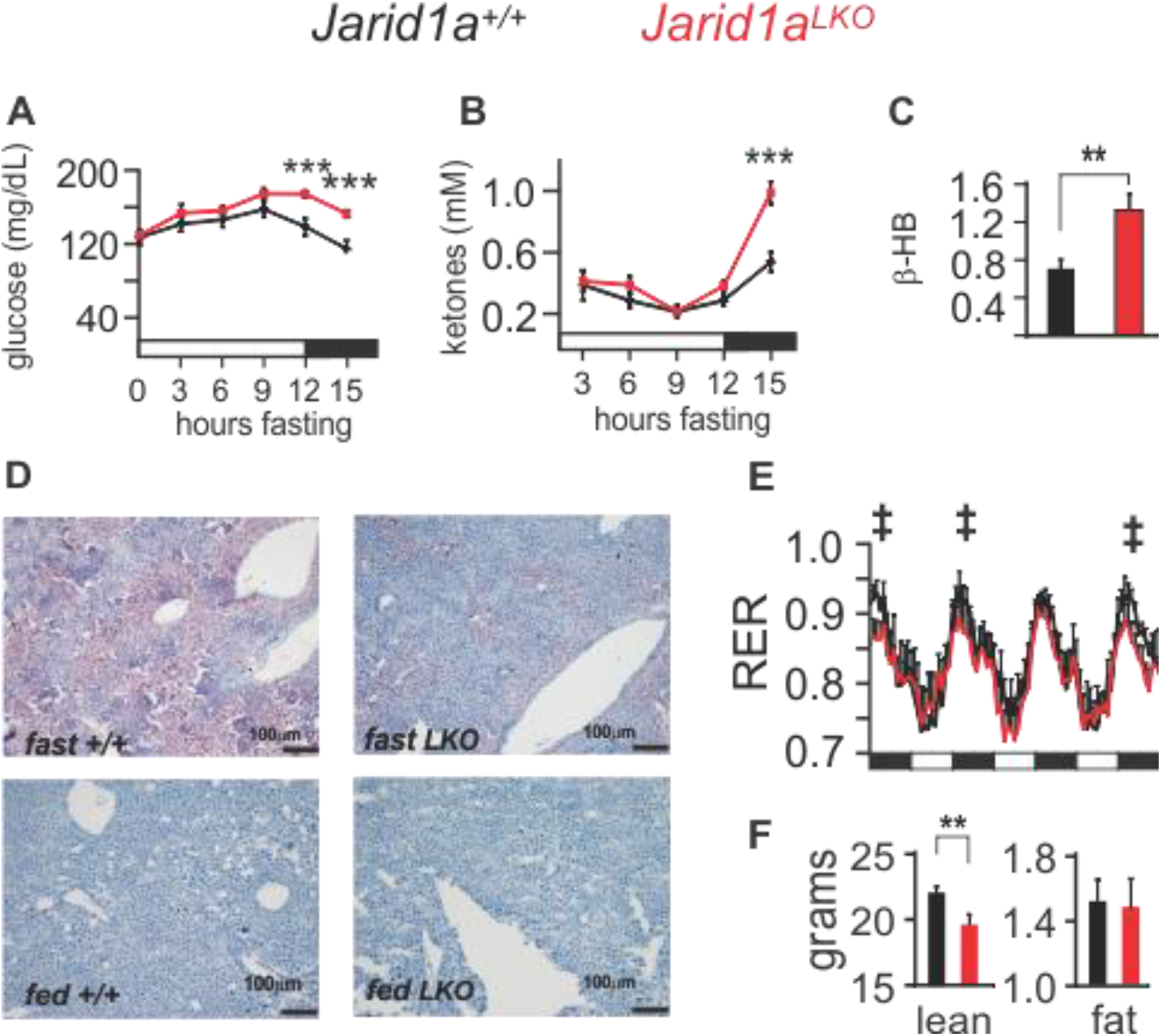
Hepatic ablation of JARID1a alters systemic glucose and lipid metabolism. (**A**) Blood glucose (***p=0.0049 and 0.0047) and (**B**) ketone levels (*** p= 0.0022) of mice subjected to fasting (mean+sem, n=8). (**C**) Hepatic β-hydroxybutyrate levels, of control and Jarid1aLKO mice (mmol/mg of tissue, mean+/− s.e.m., ** p<0.001, fig. S3 shows the time-resolved individual values). (**D**) Representative Oil Red-O staining of liver sections from fasted and fed control and *Jarid1a^LKO^* animals at 10X magnification. (**E**) Respiratory exchange ratio (‡, p<0.005 for measurements obtained from 18:00-21:00 hh, n=10). (**F**) Total lean and fat body content from control and JARID1a-null animals. Statistical data determined with a one-tailed permutation test. Black bars, dark-light cycle (dark starts at 18:00 hh, light at 6:00 hh).

However, the blood glucose of *Jarid1a^LKO^* mice remained elevated even after fasting for 15 hours, in contrast to control mice which showed a decline in blood glucose at 12 h, with a further decrease at 15 hours without food (Figure 2A). Blood ketones were undetectable in both cohorts until 3 hours after food withdrawal and remained unaltered for the next 9 hours. However, although fasting for 15 hours led to an increase in blood ketone levels in both groups, the concentration was much higher in *Jarid1a^LKO^* mice (Figure 2B). Ketones are produced by the liver following the catabolism of fatty acids, which, during a fast, are mobilized from fat stores throughout the body. In ad-lib fed animals, we found elevated average ketone levels in hepatic tissue from *Jarid1a^LKO^* mice (Figure 2C and S3). Following a 24-hour fast, wild type mice exhibit hepatic steatosis as the early influx of fatty acids exceeds the normal capacity for oxidation. In contrast, we observed protection from steatosis following a 24-hour fast in *Jarid1a^LKO^* mice (Figure 2D). Altogether, these results suggest that livers lacking JARID1a have increased capacity to provide alternative energy sources to the body during times of fasting as well as increased basal lipolysis within the liver.

We next measured respiratory exchange ratio (RER) in control and *Jarid1a^LKO^* mice across the circadian cycle. Mice have a higher RER during their active, feeding phase, which is indicative of greater use of carbohydrates for fuel. In the inactive phase, the RER drops as mice rely on fat stores, and rises again parallel to food intake as carbohydrates become more plentiful. In *Jarid1a^LKO^* mice we found that a shift towards utilization of fatty acids exists during the active phase (Figure 2E and S4). Although the liver is one of the four organs with the highest metabolic rates (along with heart, kidney, and brain) (Kinney et al., 1992), it was nonetheless striking to find that ablation of hepatic JARID1a resulted in a detectable change in RER, given that neither activity nor feeding patterns are changed (Figure 1A-D and S4) and no genetic defect occurs outside the liver. We then performed body composition analysis using Echo-MRI and discovered that the fat mass of *Jarid1a^LKO^* mice remained unaltered, but that, surprisingly, total lean body mass was significantly decreased (control, 22.27 +/− 0.42 grams; *Jarid1a^LKO^,* 19.57 +/− 0.8 grams, mean+/− s.e.m.) (Figure 2F). As muscle is the major source of amino acids used for hepatic glucose production it is possible that hepatic dysregulation of energy metabolism results in increased muscle catabolism and loss of lean mass or diversion of dietary protein to the liver at the expense of availability to the muscle. Overall, these results demonstrate that JARID1a plays a role in the regulation of hepatic energy metabolism, and its dysfunction has systemic consequences.

To better understand the basis for the *Jarid1a^LKO^* phenotype, we investigated the impact of JARID1a loss on the hepatic transcriptome via RNA-seq. We generated a circadian mRNA timeline from livers harvested every 3 hours (n=3 mice/timepoint/genotype) from either wild-type or JARID1a liver-specific null animals. We next used the JTK_CYCLE3 (Hughes et al., 2010) algorithm to identify circadian transcripts in both control and JARID1a-null datasets which we then compared. With this analysis we identified 222 genes that lost or dampened their circadian rhythmicity to an extent no longer computationally recognized as rhythmic in the livers of *Jarid1a^LKO^* mice (Group 1). Another 194 genes gained rhythmicity (Group 2) and 195 genes were non-oscillators that had altered levels of expression (>= 30% change, Group 3) (Figure 3A-C).

**Figure 3.**
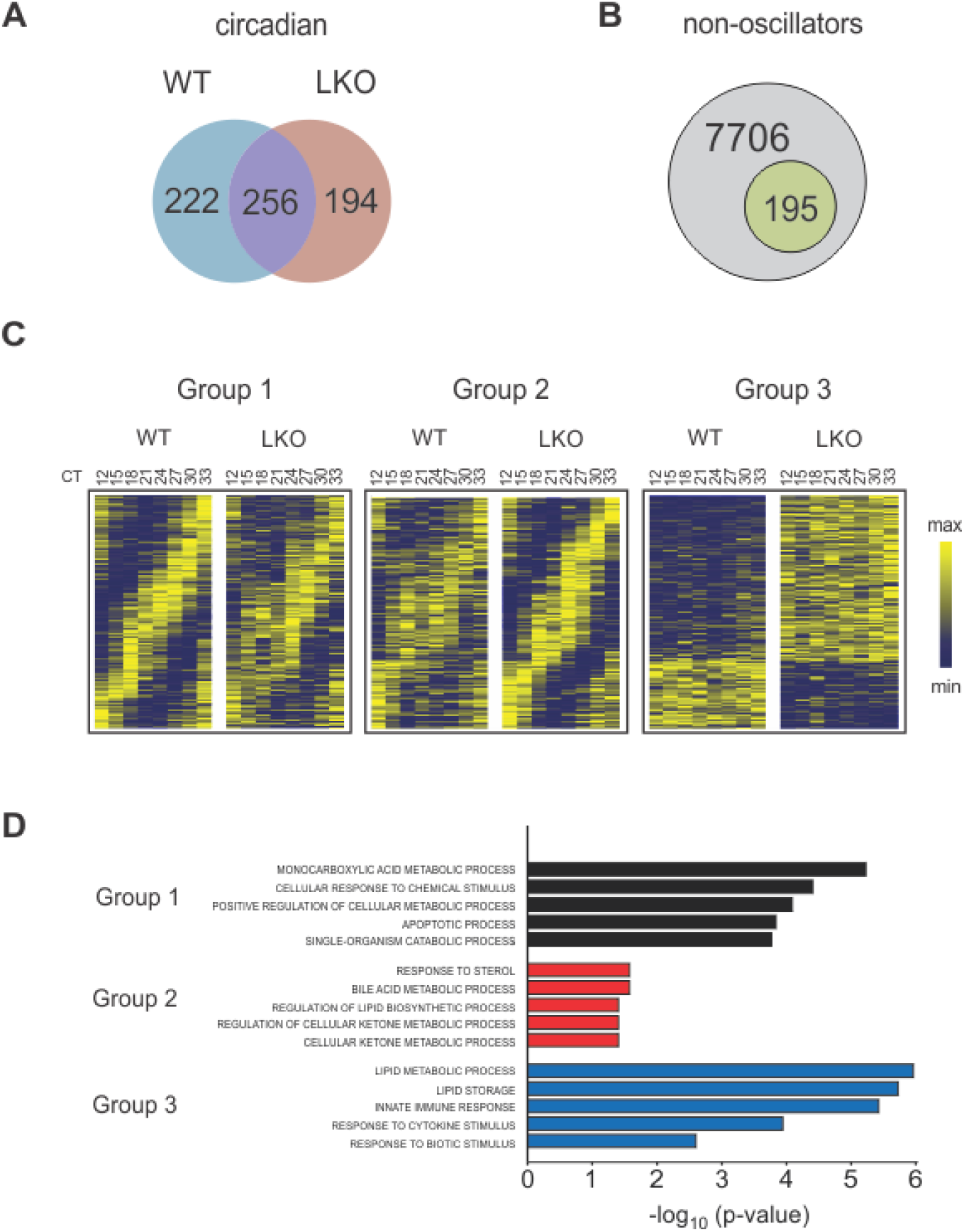
JARID1a deficiency perturbs the hepatic transcriptome. Venn diagrams of (**A**) differences and commonalities between of the circadian transcripts of control and JARID1a-null (LKO) livers, and (**B**) the number of genes scored as non-oscillating that exhibit changes to their expression levels. (**C**) Heatmaps of hepatic gene transcripts scored as rhythmic unique to control (Group 1), rhythmic unique to *Jarid1a^LKO^* (Group 2), and differentially-expressed noncircadian (Group 3). (**D**) Top five enriched ontological terms in Group1 (black bars), Group 2 (red bars), and Group 3 (blue bars).

For each group we performed ontological analysis using the DAVID bioinformatics resource (Huang da et al., 2009a, b) followed by reduction of redundant GO terms with REVIGO (Hughes et al., 2010) (Fig. 3D and Table S7). In each case, the top terms of this analysis included processes related to energy metabolism. For example, the top term associated with Group 1 was “monocarboxyl¡c acid metabolic process (G0:0032787)” whose daughter terms include “fatty acid metabolic process (GO:0006631)” and “l¡p¡d catabolic process (G0:0016042)”. Similarly, the top four terms identified in Group 2 are enriched for metabolic processes: “response to sterol (G0:0036314),” “bile acid metabolic process (G0:0008206),” and lipid and ketone metabolism (GO:0046890, GO:0010565). In the affected non-cycler group, the top parental term was oxidation-reduction process (GO:005514), whose daughter terms included processes related to carboxylic acid and lipid metabolism (GO:0019752, GO:0006629, and GO:004425).

Figure 4 shows genes affected in *Jarid1a^LKO^* mouse livers, which include mRNAs of key pro-gluconeogenic genes, including *Pck2,* whose gene product catalyzes the rate-limiting step in gluconeogenesis; *Pgc1a,* a master regulator of energy metabolism important for induction of gluconeogenesis; and *G6pc,* which allows release of glucose into the bloodstream following glycogenolysis or gluconeogenesis. Similarly, JARID1a-null livers exhibited alteration in the expression of genes linked to protein and amino acid catabolism, including elevation of components of the malate-aspartate shuttle (*Got1*), the urea cycle (*Cps1*), glutaminolysis (*Gls2*), and autophagy (*Atg101, Atg14, Atg7*). We also found reduction in the expression of lipogenic gene, *Pparg* and induction of genes involved in ketogenesis (*Bdh1, Bdh2*). We noted other disruptions to metabolic genes including shifts in peak expression, such as *Glud1* (glutamate metabolism and TCA cycle), as well as increased levels and secondary peaks of expression, such as seen in *Acmsd* (tryptophan metabolism and NAD production) and *Nnmt* (nicotinamide and xenobiotic metabolism).

**Figure 4.**
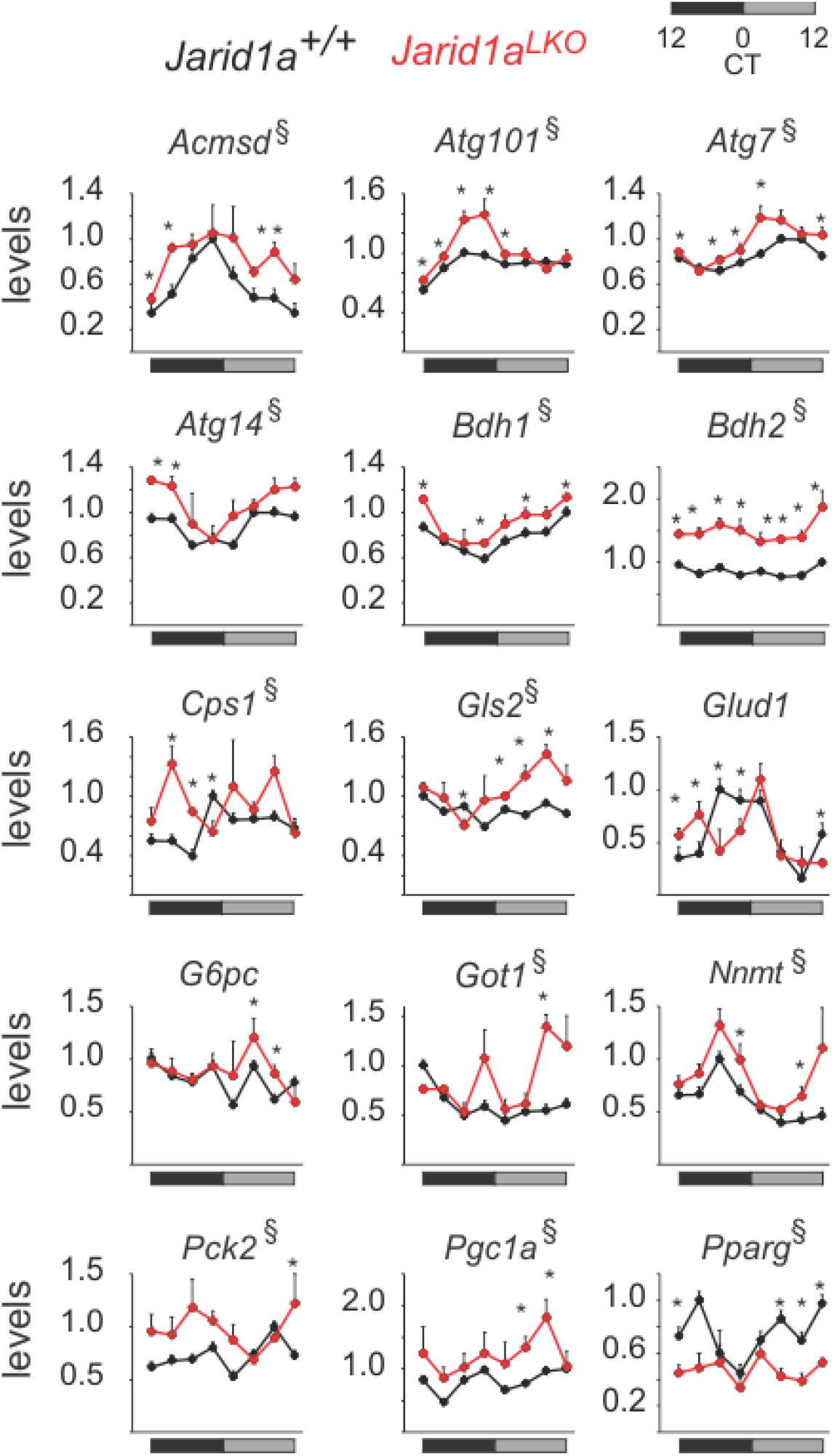
Altered energy metabolic gene expression in *Jarid1a^LKO^* livers. (**A**) Time-resolved examples of genes impacted by JARID1a ablation (3h resolution). Levels correspond to cpm normalized to the maximum value of the indicated gene in control livers (n=3, mean+sem, * denotes timepoints where p<0.05 and § where overall expression was affected with p<0.05. p-values were obtained by one-tailed permutation test.

Collectively, the gene expression data indicate that loss of *Jarid1a* disrupts metabolic gene expression. Notably, the changes we observed in *Jarid1a^LKO^* livers were reminiscent of those observed in fasted mouse liver (Sokolovic et al., 2008), in whose transcriptome there is increased gluconeogenic, lipolytic, and proteolytic gene expression. Since many metabolic genes are responsive to nutrient status, we wondered if *Jarid1a^LKO^* mice were presenting a fasted-like phenotype due to an inability to respond to nutrient availability. To test this possibility, we subjected mice to a fasted-refeeding paradigm (Figure 5A). After the last fasted phase, one cohort of each genotype was maintained under fasted conditions, and another was given access to food for 2 hours. RNA derived from mouse livers of every cohort (n=5 animals/genotype/feeding status) was sequenced and analyzed. Out of 410 food-responsive hepatic transcripts we identified in wild-type livers, 69 were dysregulated in the absence of JARID1a (Tables S10-S12). As expected, ontological analysis of this list of genes revealed enrichment of primary metabolism related terms (Figure 5B). The set included genes coding for key regulators of carbohydrate metabolism (*G6pc, Gck, Man1a*), lipid metabolism (*Apoa4, Elovl3, Lpin2*), mitochondrial function and TCA cycle (*Glud1, Coq2, Slc25a25*), and amino acid transport and protein degradation (*Dcun1d5, Slc22a28, Slc38a2, Psmd10, Rnpep, Ube2e3, Ubqln1*) (Figure 5C). Interestingly, the downregulation of several genes related to cellular stress responses (*Btg2, Cystm1, Dusp1, Ppp1r15a, Sgk1*) following feeding was defective in *Jarid1a^LKO^* livers. This defect suggests that relief from the stress of fasting is compromised in JARID1a-null liver, further supporting the idea that JARID1a plays a critical role in the fasting-feeding transition.

**Figure 5.**
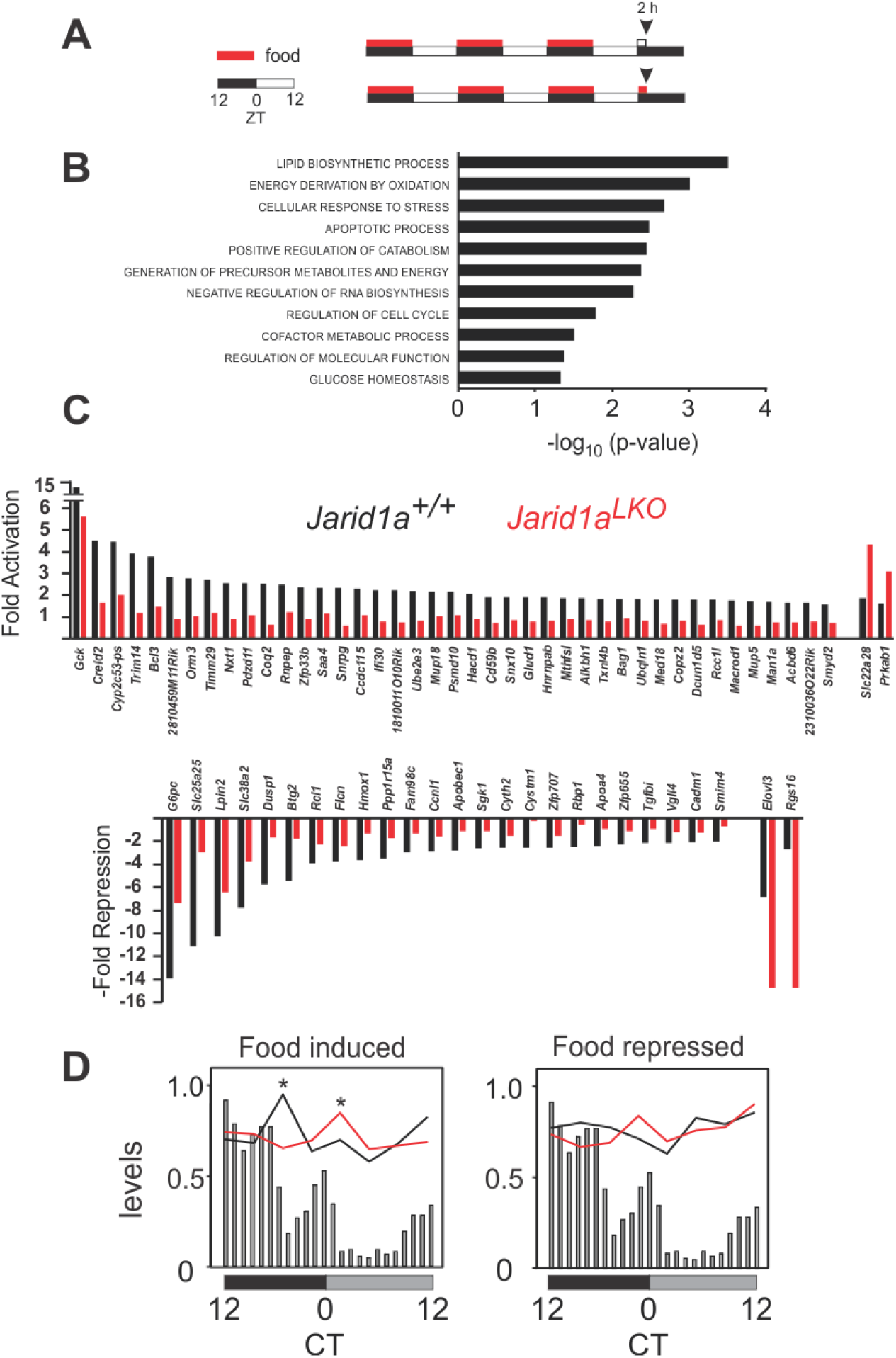
JARID1a-null livers show defective response to nutrient availability. **(A)** Schema of the experimental protocol used to assess the hepatic transcr¡ptome response to feeding. **(B)** Enriched ontological terms of food-responsive genes affected in *Jarid1a^LKO^* livers. **(C)** Food-induced (top) and repressed (bottom) genes impacted by absence of Jarid1a. Shown is fold activation from baseline levels following food consumption. Repression changes are depicted as their negative fold for ease of visualization. Specific values and statistical analysis are presented in Table S12. **(D)** Profile of food-responsive genes affected in *Jarid1a^LKO^* livers in control and JARID1a-null circadian timeline. Levels correspond to median of normalized expression levels superimposed on the circadian food intake data presented in Figure 1D (gray bars) *CT18 p-value < 0.0001 n=36; CT24 p= 0.0028, n=24; one-tailed permutation test.

Finally, we assessed the circadian profile of the food-responsive genes that were identified as affected in *Jarid1a^LKO^* livers. We grouped genes by whether they were food-induced or food-repressed in the wildtype, and tracked the expression of these subsets in our circadian RNA-seq dataset. A clear pattern emerged in which the expression of the food-induced gene set was disrupted (depicted as the median of normalized values for each gene set at each timepoint overlaid on time-resolved feeding data, see Table S13 and Methods section). Specifically, the rise in expression seen in control livers following the major feeding bout was absent in the *Jarid1a^LKO^* livers. Instead, overall levels were offset ~6h, so that a maximum was observed only following the secondary feeding session (Figure 5D). These results indicate that JARID1a is part of the mechanism by which feeding-derived cues induce the transcriptional response needed for the liver to adapt to exogenous nutrient availability.

## Discussion

The liver is the major metabolic organ of the body, responsible for coordinating relevant responses to both predictable fluctuations and acute changes in energy availability and demand. In mammals, glycolysis, lipogenesis, and global protein synthesis generally coincide with the active period, when feeding takes place. Conversely, during the inactive, fasting period, energy homeostasis is maintained via the use of internal energy stores generated during the feeding period, as glycogenolysis, lipolysis, ketogenesis and proteolysis. However, the timing of these different processes is not just a broad temporal segregation to the active-inactive period. Carbohydrate intake is preferred in the early active phase, with a switch in preference for protein and lipid towards the end of the active period (Tempel et al., 1989). Consistently, in numerous species including humans, glycolysis and lipogenesis peak in the early active phase, while glycogenesis peaks in the late active phase (Doi et al., 2010; Ishikawa and Shimazu, 1980; Panda et al., 2002). Conversely, glycogenolysis occurs during the early part of the inactive phase, while lipolysis and gluconeogenesis peak at the end of the inactive phase, as glycogen stores decline. Also coinciding with this time, the process of autophagy exhibits a circadian rhythm with maximal function at the end of the inactive period (Ma et al., 2011) when preferred energy stores have been exhausted, thus providing amino acids for energy or glucose production.

In the liver, temporal orchestration of metabolism is accomplished in large part via the imposition of large-scale oscillations in transcription, including genes that code for regulators of rate-limiting steps of anabolic and catabolic pathways (Panda et al., 2002). Here we find that *Jarid1a* ablation results in noticeable alterations in the expression profile of regulators of metabolism, including overall levels and/or circadian patterns. Although important, the hepatic clock by itself only partly accounts for the large-scale transcriptional rhythms observed in the liver. In synergy with the clock, feeding cues drive a large component of oscillatory genes and also feedback to entrain the clock (Schibler et al., 2003; Vollmers et al., 2009). This synergistic system allows the liver to anticipate the predictable fluctuating energy demands of the body while also maintaining the ability to respond to acute feeding cues. Our data implicates JARID1a in this circadian clock-feeding interaction, and suggests that its presence is important in the management of nutrient availability.

The liver impacts energy homeostasis for the whole body and the effects of hepatic JARID1a deficiency can be observed at that scale. First, *Jarid1a^LKO^* animals experience less weight gain over time-an effect that is more pronounced when the mice are given a high-fat diet. Second, they exhibit lower respiratory exchange ratios (RER), indicative of increased fat oxidation. Finally, *Jarid1a^LKO^* animals exhibit decreased total lean body mass. At the tissue level, we found that loss of JARID1a led to disruption of metabolic gene expression. *Jarid1a^LKO^* livers were defective in their transcriptional response to feeding challenge and demonstrated an overall profile similar to a fasted state. Altogether, our results show that the clock component JARID1a has an important role in hepatic adaptation to exogenous nutrient availability, with critical relevance for both local and systemic metabolic health.

## Author contributions

K.D. designed study, carried out the research, analyzed and interpreted results, and wrote the manuscript. D.K. carried out the research and analyzed data. A.S. performed experiments. A.B. performed experiments. C.V. performed experiments, analyzed and interpreted data, and reviewed the manuscript. L.D. designed the study, carried out research, analyzed and interpreted results, wrote and reviewed the manuscript, and is responsible for the integrity of the work.

## Acknowledgments

We would like to thank Shubhroz Gill and Partha Kasturi for technical advice and comments on the manuscript; the KUMC Laboratory Animal Resources, Metabolic Core, and the Department of Pharmacology Core facilities for technical assistance. Mice strains are commercially available from Jackson Laboratory. The data presented in this report is found either in the main text or in the supplementary materials. RNA sequencing data have been made available at NCBI’s GEO. This work was supported by grants from the National Institute of Diabetes and Digestive and Kidney Diseases under grant number R01DK108088 (to LD), National Center for Research Resources (P20RR021940) and the National Institute of General Medical Sciences (P20GM103549) of the National Institutes of Health, Institutional Development Award (IDeA) from the National Institute of General Medical Sciences of the National Institutes of Health under grant number P20 GM103418, and the Lied Basic Science Program (to LD.)

## Materials and Methods

### Mice

Liver-specific JARID1a-null (animals were generated by breeding *Jarid1a^loxP/loxP^* mice (C57BL/6J congenic, Jackson Laboratories stock 008572) with a transgenic mouse line that specifically expresses CRE in the liver at high levels (Albumin promoter-driven CRE; C57BL/6J congenic Alb-CRE, Jackson Laboratories stock 003574. LoxP(-)/Cre+ littermates were used as controls.

### Body weight measurements

8-week-old Jarid1a^LKO^ and control mice were housed under a 14 h: 10 h light dark cycle with access to either chow or a high-fat diet (40% kcal from fat). Mice were subjected to weekly whole body weight measurements for 18 weeks.

### Glucose and Pyruvate tolerance tests

Mice were fasted for 16 h prior to testing. After this period, the lateral tail vein was incised to allow for blood collection. Fasted blood glucose was measured (FreeStyle Lite) before injection and recorded. Each mouse was injected with either pyruvate (2 mg/g body weight) or glucose (1mg/g body weight). Subsequent measurements were made by collecting blood from the cut site before injection and at 30, 60, and 120 minutes after injection.

### Activity recordings

*Jarid1a^LKO^* mice were maintained under a 12 h:12 h LD cycle. Total activity of mice housed in a light-tight, custom-made circadian chamber was recorded and analyzed with a BigBrother video monitoring system (Figure 1A and S1). In addition, activity was also analyzed by beam break recordings during the metabolic cage study (Figure 2D) (Promethion system, Sable).

### Food consumption

Following entrainment to a 12 h:12 h LD cycle in a light-tight chamber, individually housed mice were transferred to a Minimitter chamber (Colombus Instruments). Total daily food consumption and meal pattern analysis of mice under 12 h:12 h LD was done for three days (Figure 1C). In addition, total food consumption and meal pattern analysis was independently done as part of the metabolic cage study Figure S4) (Promethion system, Sable).

### b-Hydroxybutyrate measurements

Assays were performed with Cayman Chemical’s beta-Hydroxybutyrate Colorimetric Assay kit (Cat#700190).

### Histology

Mice were either fasted for 24 h or maintained with ad libitum access to food, sacrificed by cervical dislocation, and necropsied. For each cohort four different mice were used. Whole body weight was recorded and liver tissue was collected. Liver sections were cryopreserved, sectioned with a cryostat, and stained with Oil red O.

### Metabolic cages

Following entrainment to a 12 h:12 h LD cycle in a light-tight chamber, individually housed mice were transferred to a Promethion metabolic cage system (Sable Systems International, Las Vegas, NV). Recordings of respiratory exchange ratio, VO2, activity, and food consumption of 10-14 week old animals was done for a total of 5 days under a 12 h: 12 h LD cycle.

### Body composition

Total lean and fat mass analysis was done in an EchoMRI-1100 (EchoMRI LLC, Houston, TX) instrument following the manufacturer’s instructions.

### Circadian tissue timeline generation

Mice were housed in a homemade light-tight chamber where they were entrained to a 12 h: 12h light-dark cycle for two weeks. Following entrainment mice were released into total darkness for 24 hours at which point they were sacrificed by cervical dislocation and necropsied at 3 hour intervals. For each genotype an n=3 per timepoint were collected.

### Fasting blood glucose and ketones

To reduce intra-sample variability arising from naturally-occurring differences in consumption patterns in different mice, mice were subjected to 12 h fasting: 12 h feeding paradigm for three days. At the end of the third feeding period, food was removed and blood glucose and ketone in fasted mice were respectively monitored with a FreeStyle Lite blood glucose and with a NovaMax Plus Ketone meters.

### Fasted-refeeding tissue harvesting

Mice were subjected to a three-day 12 h fasting: 12 h feeding paradigm. At the end of the third fasting period mice of each genotype were either maintained under fasting conditons or allowed access to food for two hours before being killed (Figure 5A). Each cohort consisted of n=5. RNAseq Library Preparation RNA was extracted from control and *Jarid1a^LKO^* liver samples using trizol extraction. RNAseq libraries were then constructed using a variation of the Smartseq2 method (Picelli et al., 2013). In short, we generated cDNA from total RNA using the Smartscribe Enzyme (Clontech) and a oligo-dT primer modified with a universal priming site (ISPCR). During reverse transcription another universal priming site (ISPCR) was added at to the end of the cDNA corresponding to the 5’ end of transcripts through template switching to an additional oligo. The resulting full-length cDNA was treated with RNAse A and Lambda Exonuclease to remove template-switch oligos and primer dimers, respectively and amplified using the universal priming sites with KAPA HIFI Hot Start Readymix (KAPA) and a ISPCR primer. The PCR product was then treated with Tn5 enzyme loaded with Tn5ME A/R and Tn5ME B/R adapters. The resulting fragments were amplified using Nextera A and Nextera B index primers using KAPA HIFI DNA Polymerase (KAPA), size selected to 350-800bp, and sequenced on a Illumina HiSeq 2500.

Nextera_A_Index_Primer,

AATGATACGGCGACCACCGAGATCTACAC[i5]TCGTCGGCAGCGTCAGATG

Nextera_B_Index_Primer,

CAAGCAGAAGACGGCATACGAGAT[i7]GTGGGCTCGGAGATGTGTAT

Tn5ME-R, [phos]CTGTCTCTTATACACATCT

Tn5ME-A (Illumina FC-121-1030), TCGTCGGCAGCGTCAGATGTGTATAAGAGACAG

Tn5ME-B (Illumina FC-121-1031), GTCTCGTGGGCTCGGAGATGTGTATAAGAGACAG

Smartseq2-TSO, AAGCAGTGGTATCAACGCAGAGTACATrGrGrG

OligodT/iMeCisodC/AAGCAGTGGTATCAACGCAGAGTACT30VN

ISPCR, AAGCAGTGGTATCAACGCAGAGT

### RNA-seq Analysis

Circadian timeline: Fastq files from the Illumina HiSeq2500 run aligned to GRCm39 version of the mouse genome using the STAR aligner (Dobin et al., 2013). Raw counts were normalized to counts per million (CPM), and the cutoff of expression was defined as CPM >= 1 in 41 out of the 48 libraries sequenced. Expressed genes were analyzed with JTK_CYCLE3, and rhythmic genes were defined on the basis of relative amplitude >0.2 and FDR<0.1. Non-circadian genes were considered to be differentially expressed if they exhibited a 30% change and FDR<0.1. Heatmaps were generated with the Broad Institute’s Morpheus tool (https://software.broadinstitute.org/morpheus/). Fasting-refeeding libraries were sequenced in an Illumina HiSeq2500 run. The resulting Fastq files were aligned to GCRm39 version with Cufflinks and expressed genes defined as those with for which RPKM >0.5 in 17 out of the 20 libraries sequenced. For differential expression analysis, a cutoff of 50% change and FDR<0.1 were used. One-tailed p-values were calculated with a permutation test (5000 Monte Carlo simulations), which were then used to calculate the FDR via the Benjamini-Hochberg correction.

---

**Fig. S1.**
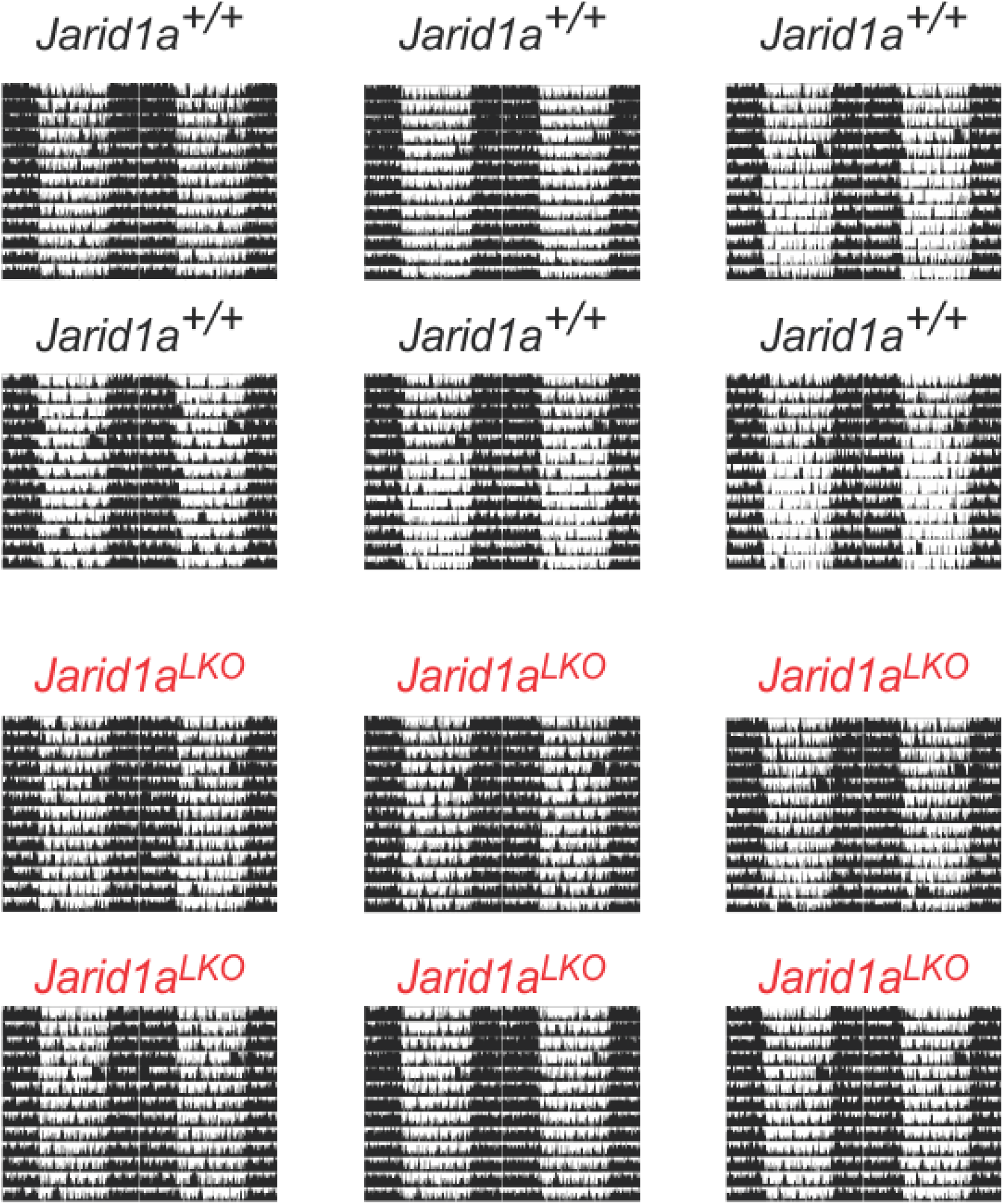
Double-plotted actogram recordings of control and Jarid1a^LKO^ animals housed in a circadian light-tight chamber under a 12 h: 12h LD cycle.

**Fig. S2.**
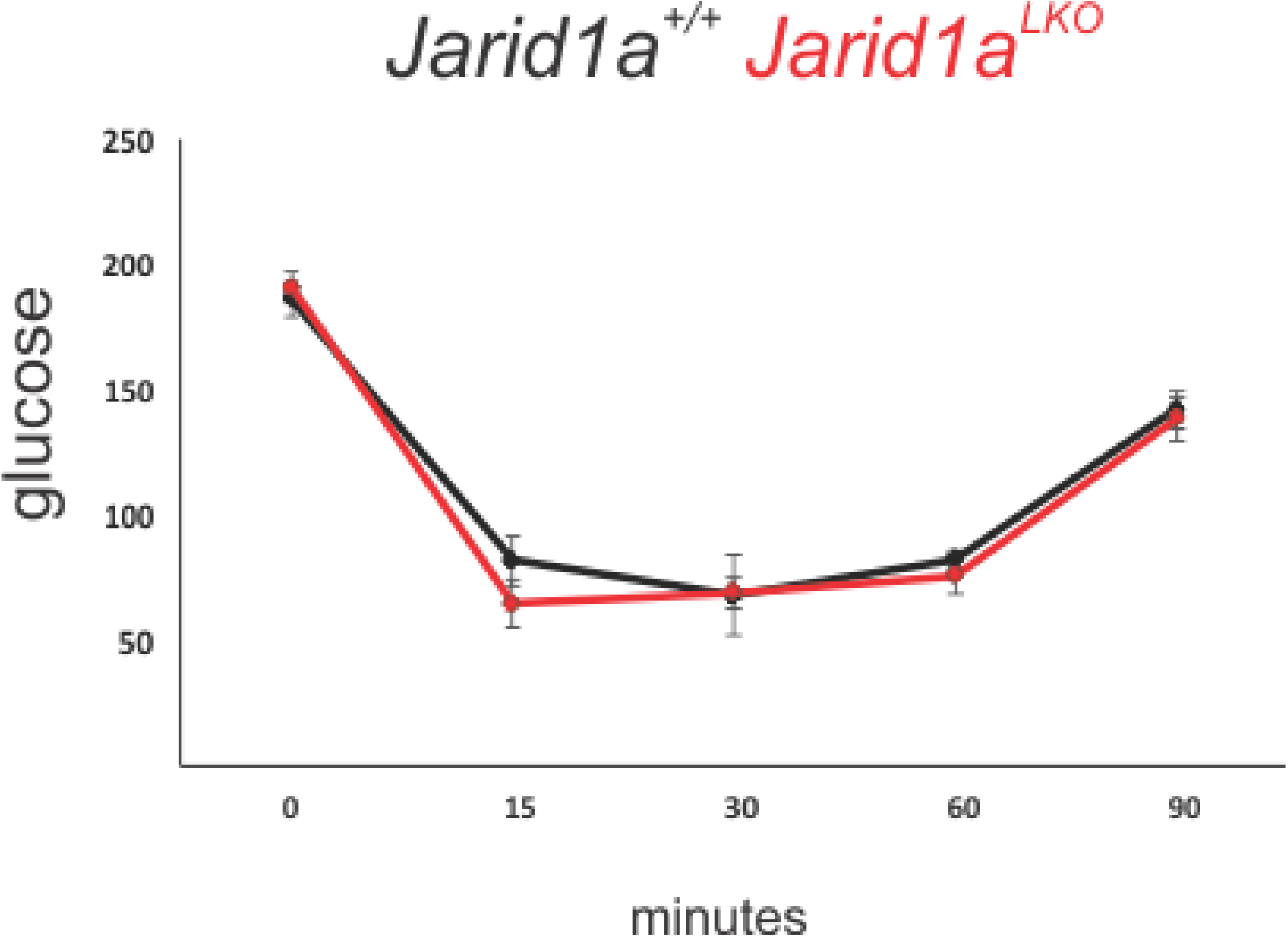
Insulin tolerance test of control and Jarid1aLKO mice

**Fig. S3.**
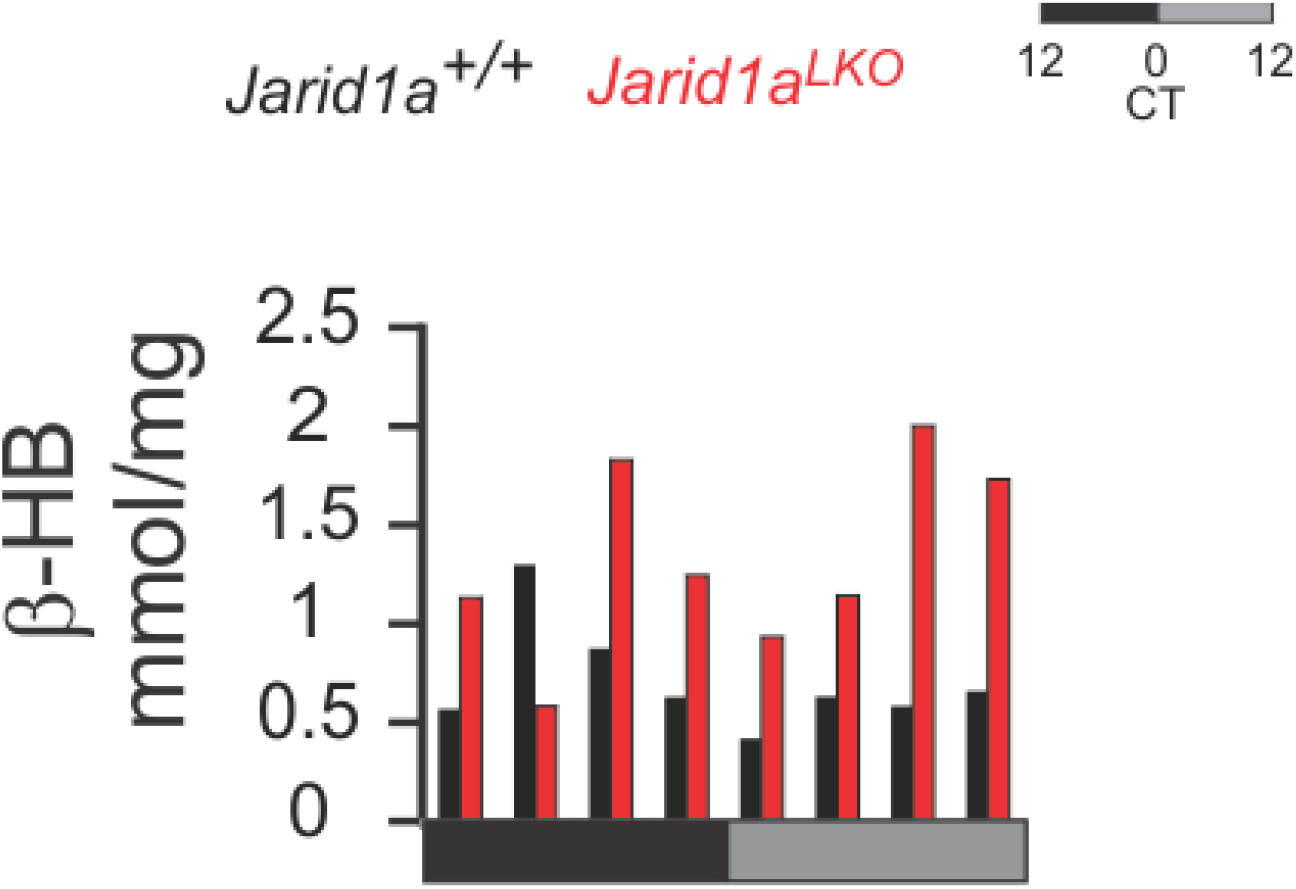
Time-resolved b-HB levels shown in Figure 2C.

**Fig. S4.**
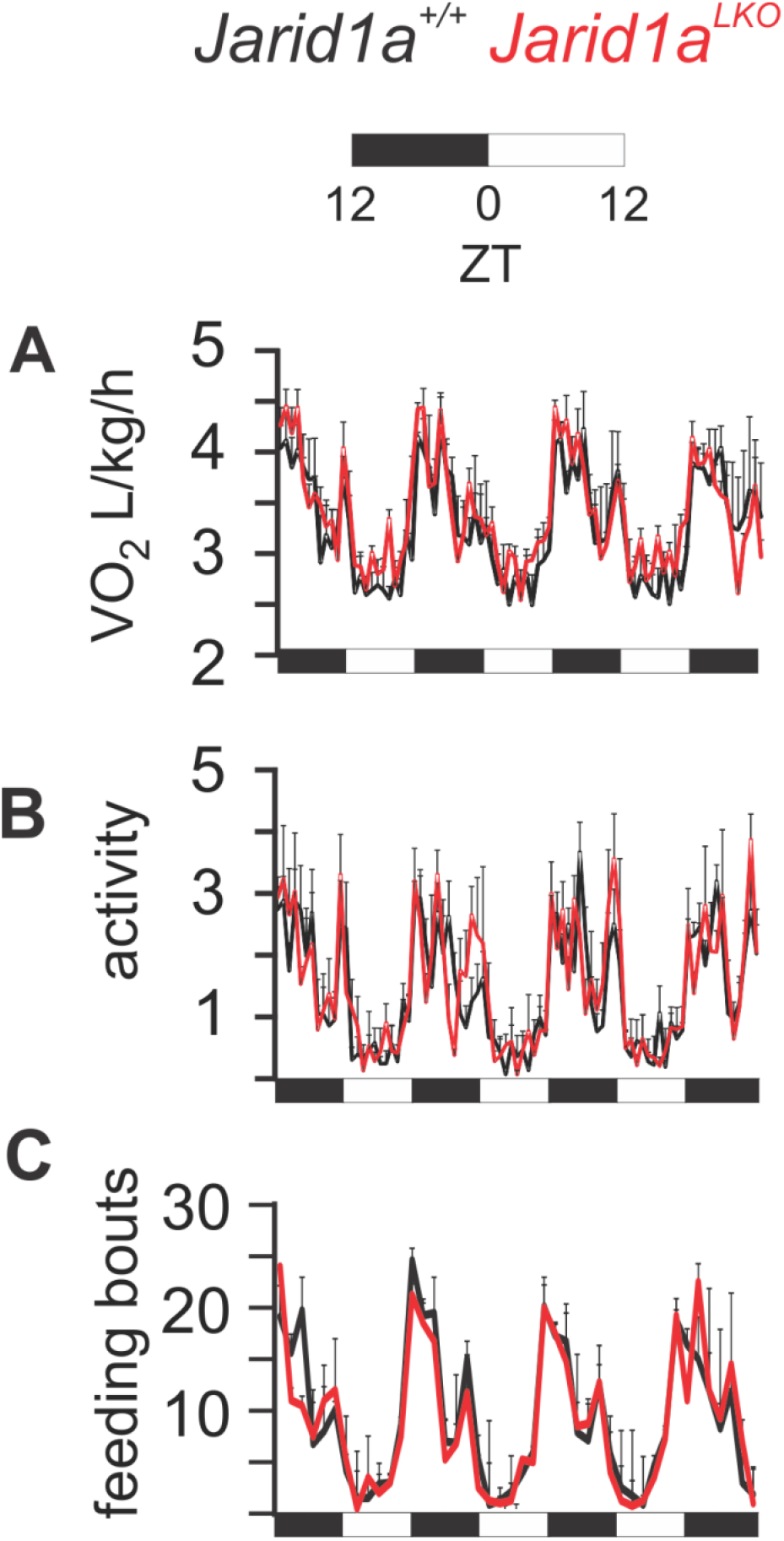
(A) O2 consumption normalized to body weight, (B) activity measured as the sum of Y-axis and X-axis beam breaks per hour (x 10^3^), and (C) Feeding bouts per hour, collected in simultaneous to Figure 3C.

## Supplementary Table legends

**Table S1. Expressed genes in circadian timelines.**

**Table S2. Genes scored as circadian in control and arrhythmic in JARID1a-null timelines.**

**Table S3. Genes scored as circadian in JARID1a-null livers but not control timelines.**

**Table S4. Genes scored as rhythmic in both control and JARID1a-null timelines. Tables S5. Noncircadian, differentially-expressed genes.**

**Table S6. All noncircadian genes.**

**Table S7. Ontology analysis of gene sets derived from the circadian timelines.**

**Table S8. Specific p-values for Figure 4.**

**Table S9. Gene datasets for Fasted-refeeding experiments.**

**Table S10. Genes identified as food responsive in control livers.**

**Table S11. Food responsive genes impacted by *Jarid1a* ablation.**

**Table S12. Individual values and calculations pertaining to figure 5D.**

